# An engineered disulfide staple restricts lid loop dynamics and alters substrate specificity of phenylalanine ammonia-lyase

**DOI:** 10.64898/2026.05.01.722275

**Authors:** Rebecca Condruti, Likith Muthuraj, Jeevan K. Prakash, Samuel D. Littman, Pravin Kumar R., Nikhil U. Nair

## Abstract

In *Anabaena variabilis* (*Trichormus variabilis*) phenylalanine ammonia-lyase (*Av*PAL), a conserved lid-like loop sits over the active site and has been studied both for its role in positioning a catalytic tyrosine and for its contribution to phenylalanine aminomutase (PAM) activity. While the active site architecture and substrate specificity of *Av*PAL have been extensively characterized, the dynamic behavior of this unstructured loop beyond its role in catalysis remains poorly understood. Here, we investigate the functional role of this loop by restricting its mobility through targeted interchain disulfide bond engineering. Three in-house approaches were designed to predict ideal cysteine residue pairs: (i) quantifying pair interaction energies via electrostatic and van der Waals forces, (ii) generating a contact map of residues within 5 Å proximity, and (iii) implementing a machine-learning model trained on datasets from PDBCYS, SPX, and an internal database to rank cysteine pair likelihood within disulfide bond geometric constraints. Our machine-learning-guided strategy yielded a successful variant with complete oxidation efficiency in *E. coli*. Rigidification of this loop reveals that it also functions as a regulator of substrate specificity. Multi-scale molecular simulation analyses (molecular dynamics, metadynamics, quantum/molecular mechanics) reveal that this modification alters the active-site pocket by reducing the conformational dynamics of substrate binding. Our findings underscore the delicate balance between enzyme flexibility and catalytic efficiency, providing novel insights into the role of this understudied dynamic loop region in *Av*PAL.

## INTRODUCTION

Phenylalanine ammonia-lyase (PAL) has been extensively studied as a therapeutic in treating phenylketonuria (PKU), its essential role in the plant phenylpropanoid pathway, and its capacity to synthesize industrially relevant compounds (1). It catalyzes the non-oxidative deamination of L-phenylalanine to *trans*-cinnamic acid and ammonium via its unique, catalytic moiety, 4-methylidene-imidazole-5-one (MIO) (2). While named for its L-phenylalanine (phe) substrate specificity, PAL exhibits limited substrate promiscuity toward other aromatic amino acids and can also display weak aminomutase-like activity (3, 4). Multiple studies have shown that this preference is governed by a combination of conserved active-site residues and the conformational dynamics of flexible loop regions surrounding the catalytic pocket.

Despite extensive engineering efforts and biochemical characterization across the PAL family, the dynamic behavior of multiple unstructured loop segments that interact with the active site remains understudied. In *Anabaena variabilis* phenylalanine ammonia-lyase (*Av*PAL), a “lid-like” loop has been shown to influence lyase and mutase activities (**Figure 1a**). This loop is the most extensively characterized unstructured region, as it contains a conserved catalytic tyrosine that directly interacts with the MIO moiety and participates in substrate positioning. Studies have demonstrated that mutations in this region improve activity, suggesting that this loop also plays a role in steering catalytic preference (5, 6). Together, these observations suggest the lid-like loop functions as a dynamic element that integrates flexibility with a fine-tuned catalytic outcome.

**Figure 1.**
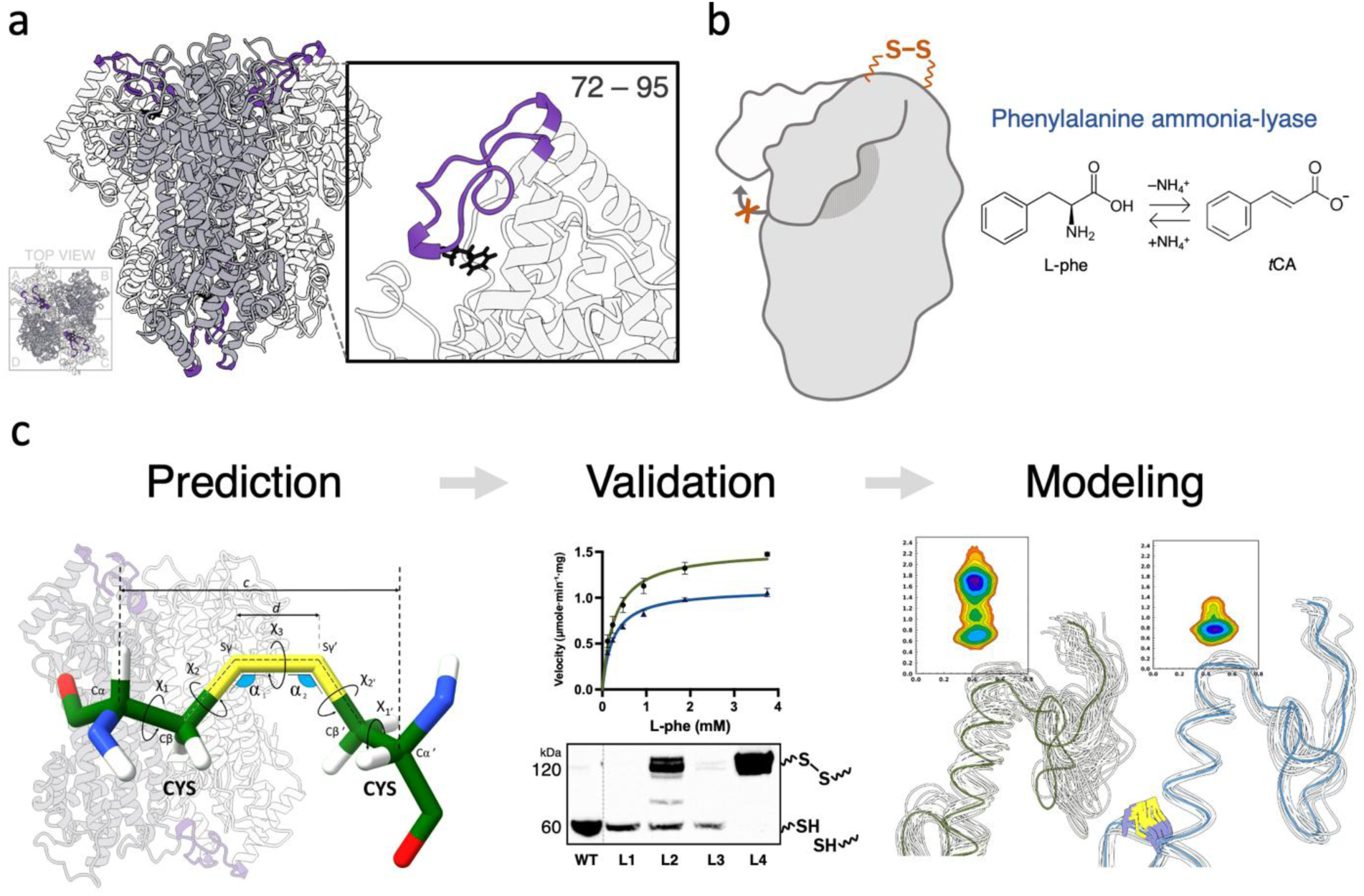
Overview of this work. (a) The lid-like loop (residues 72–95) of *Av*PAL is highlighted (purple) above the substrate (black). (b) Reaction catalyzed by *Av*PAL, showing the non-oxidative deamination of L-phenylalanine to *trans*-cinnamate with release of ammonium. (c) Computational and experimental workflow used to identify and evaluate engineered disulfide bonds in *Av*PAL.

In this study, we investigate the role of the lid-like loop in *Av*PAL by modulating its flexibility and assessing the functional consequences of restricting its conformational freedom via targeted disulfide bond engineering (**Figure 1b**). Disulfide bonds have been widely utilized in protein engineering to enhance stability and improve folding efficiency (7, 8). To guide disulfide bond placement, we built three predictive frameworks to identify cysteine pairs that formed interchain disulfide bonds (**Figure 1c**). We introduced a disulfide bond at the hinge of the loop that restricted its flexibility and substrate specificity. To complement our experimental approach, we employed molecular simulations to gain mechanistic insights into how loop dynamics influence catalytic interactions. Our results indicate that modulating the flexibility of the lid-like loop serves as a regulator of substrate specificity, with decreased flexibility reducing substrate promiscuity. Our results highlight the balance between a known flexible region in *Av*PAL and its subtle biochemical regulation, with decreased flexibility diminishing the enzyme’s ability to accommodate alternative substrates within the active-site pocket. These findings provide valuable insight into the possibility of unstructured loop regions serving as active contributors to catalysis, rather than passive structural elements. By examining the structural and functional consequences of rigidification, this study sheds light on how we may be oversimplifying and overlooking the role of flexible loops in enzymatic catalysis.

## RESULTS

### Custom framework enables targeted disulfide engineering in the lid-like loop of *Av*PAL

Numerous studies have demonstrated the utility of engineered disulfide bonds in enzymes, yet the presence of structurally permissive sites is inherently protein-specific and governed by each enzyme’s unique structure. The feasibility of both intrachain and interchain disulfide bonds is highly dependent on the protein architecture, as *de novo* cysteines must adhere to the strict set of geometric and stereochemical constraints, including appropriate Cβ:Cβ spacing, backbone orientation, dihedral angles, and solvent exposure for redox feasibility (9). Disulfide bond engineering within the lid-like loop of *Av*PAL has not been explored, to our knowledge, and the prediction of engineered disulfide bonds remains limited by the scarcity of accessible tools. This motivated the development of computational calculations to exhaustively generate all feasible disulfide bond placements within a defined structural region via distinct methods. We designed three methods to predict disulfide bonds in *Av*PAL: pair interaction energy maps (PIE), contact maps, and a machine learning (ML) algorithm (**Figure 2**). Each approach independently highlighted multiple residue pairs with favorable energetic, structural, and dynamic characteristics for disulfide bond formation. We evaluated the feasibility and stability of disulfide bond formation through molecular dynamics simulations conducted over a temperature range of 303.15 K to 373.15 K. We filtered variants based on multiple stability criteria, including low backbone RMSD (root mean square deviation), preservation of active-site flexibility as assessed by RMSF (root mean square fluctuation), and maintenance of a stable sulfur-sulfur distance of approximately 2.5 Å throughout the simulations, indicative of persistent disulfide bond formation without disruption of the active-site architecture. Based on these criteria, we shortlisted a subset of four disulfide pairs for experimental evaluation (**Table 1**). The first two methods each generated one residue pair, and the third method generated two pairs. Our methods did not predict any intrachain disulfide bonds.

**Figure 2.**
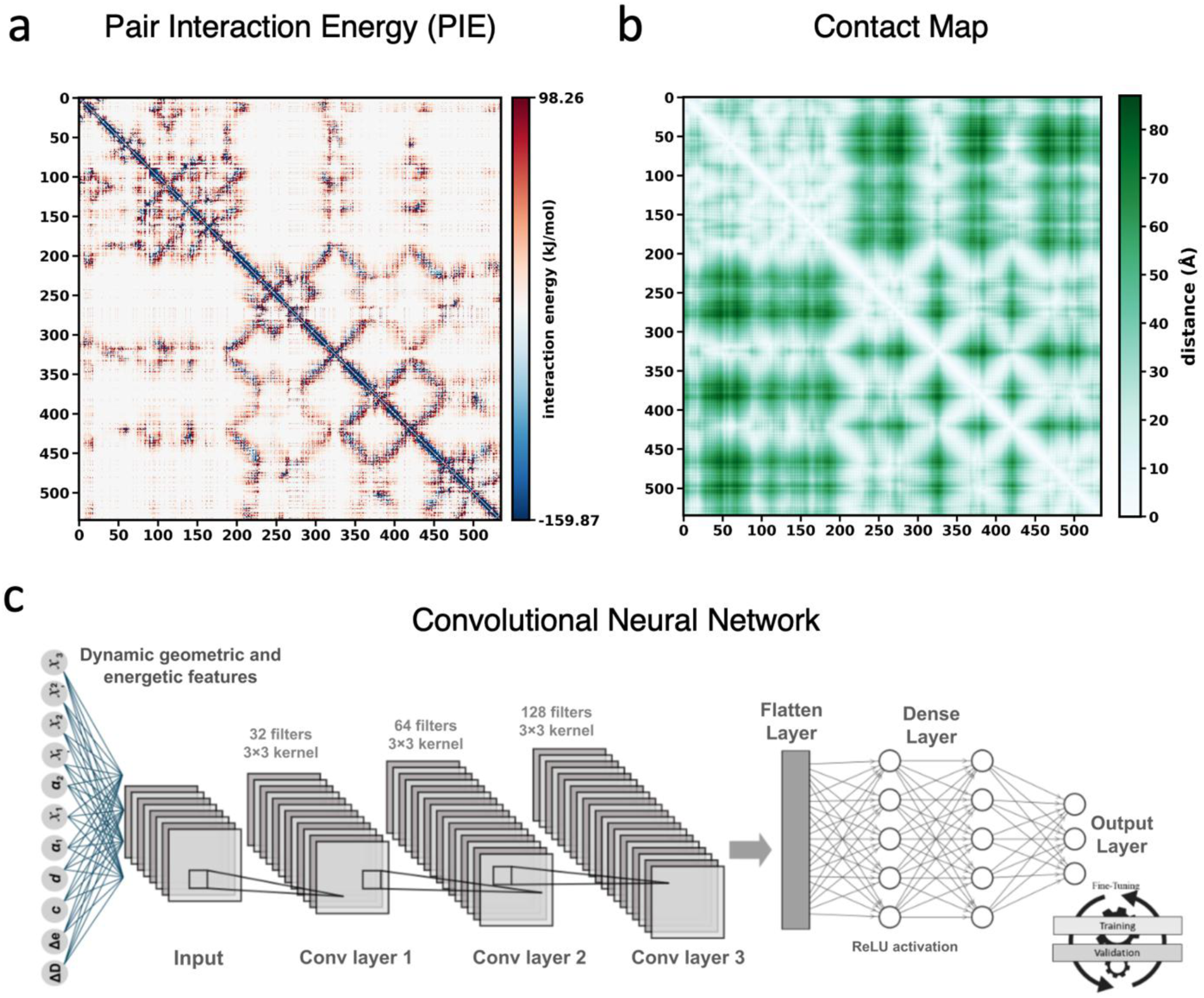
The prediction strategies for disulfide bond engineering. (a) Paired interaction energy (PIE) maps, (b) contact of residues map, and (c) a machine learning (ML) pipeline utilizing geometrical principals of disulfide bonds. The residue scores from PIE and contact maps can be found in the Supporting Data File.

**Table 1.** Prediction methods and mutations for disulfide bond formation.

| Variant | Residue 1 | Residue 2 | Pred. Method |
| --- | --- | --- | --- |
| **L1** | T110 | L421 | Contact maps |
| **L2** | F107 | N533 | ML |
| **L3** | N103 | V414 | PIE |
| **L4** | W106 | N534 | ML |

### Whole-cell activity screening identifies active *Av*PAL variants with cysteine substitutions

Since the disulfide-prediction pipeline relied solely on geometric and structural criteria, it could not predict how cysteine substitutions would influence catalytic efficiency, folding, or stability. Thus, we implemented a semi-high-throughput spectrophotometric whole-cell assay to quantify changes in enzyme activity resulting from the replacement of native residues with engineered cysteine pairs (5). Activity measurements revealed that most engineered variants exhibited reduced activity relative to WT (**Figure 3a**). Variants L1, L2, and L3 displayed substantial decreases in activity, each retaining only 40–60 % activity. In contrast, variant L4 largely retained its activity (**Figure 3b**).

**Figure 3.**
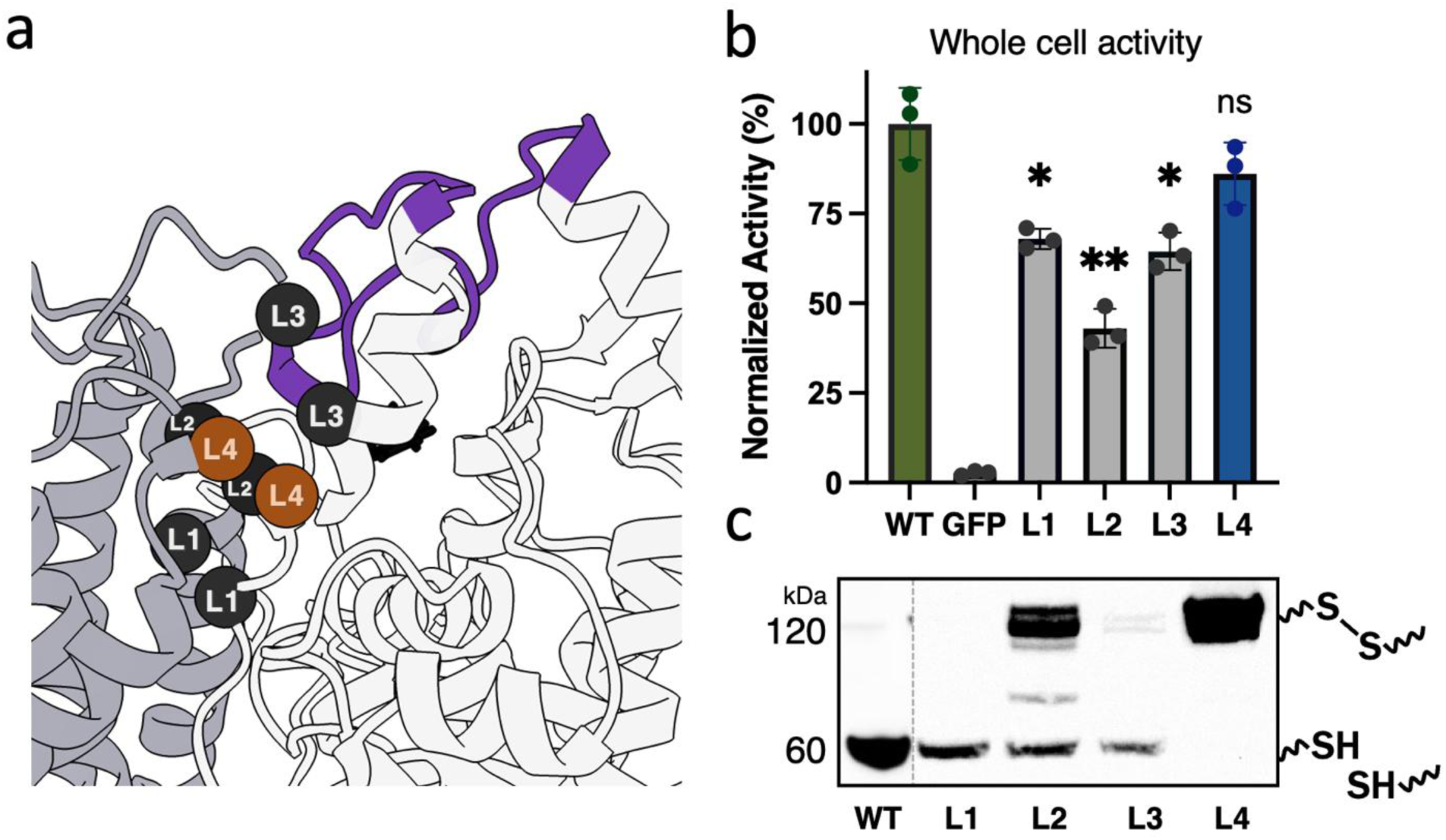
Experimental validation of disulfide bond predictions. (a) Location of interchain disulfide bonds (L1 – L4). Variant L4, the focus of this study, is highlighted in orange. (b) A whole-cell activity assay was used to determine how cysteine substitutions affected activity. (c) Non-reducing Western blot of cell lysates shows that only L4 forms complete dimers with intersubunit disulfide bonds. Two-tailed unpaired t-test, * p < 0.05, ** p < 0.005. WT = wild-type. GFP = green fluorescent protein, control.

#### Variant L4 forms a robust interchain disulfide bond within the lid-like loop of *Av*PAL

After measuring changes in activity from engineered disulfide bonds, we sought to determine whether the engineered disulfide bonds successfully formed *in vivo*. Disulfide formation requires an oxidative intracellular environment, a condition not supported in traditional *E. coli* expression strains, as they maintain a reducing cytoplasm. We therefore used *E. coli* NEB SHuffle, which has been engineered to support disulfide bonds formation. In bacterial lysates, variants L1 and L3 showed minimal evidence of disulfide-mediated dimerization; the predominant species migrated at the monomeric molecular weight, indicating that these cysteine pairs either failed to oxidize or were structurally positioned in a manner that precluded disulfide bond formation (**Figure 3c**). Variant L2 displayed both monomeric and dimeric bands, suggesting the formation of a mixed population in which only a subset of subunits successfully form the disulfide bond. In contrast, variant L4 was fully oxidized. This oxidation was visible via Western blot, where the denatured protein migrates with a molecular weight of a dimer. The oxidized state is stable and is maintained during affinity purification (**Figure 4a**).

**Figure 4.**
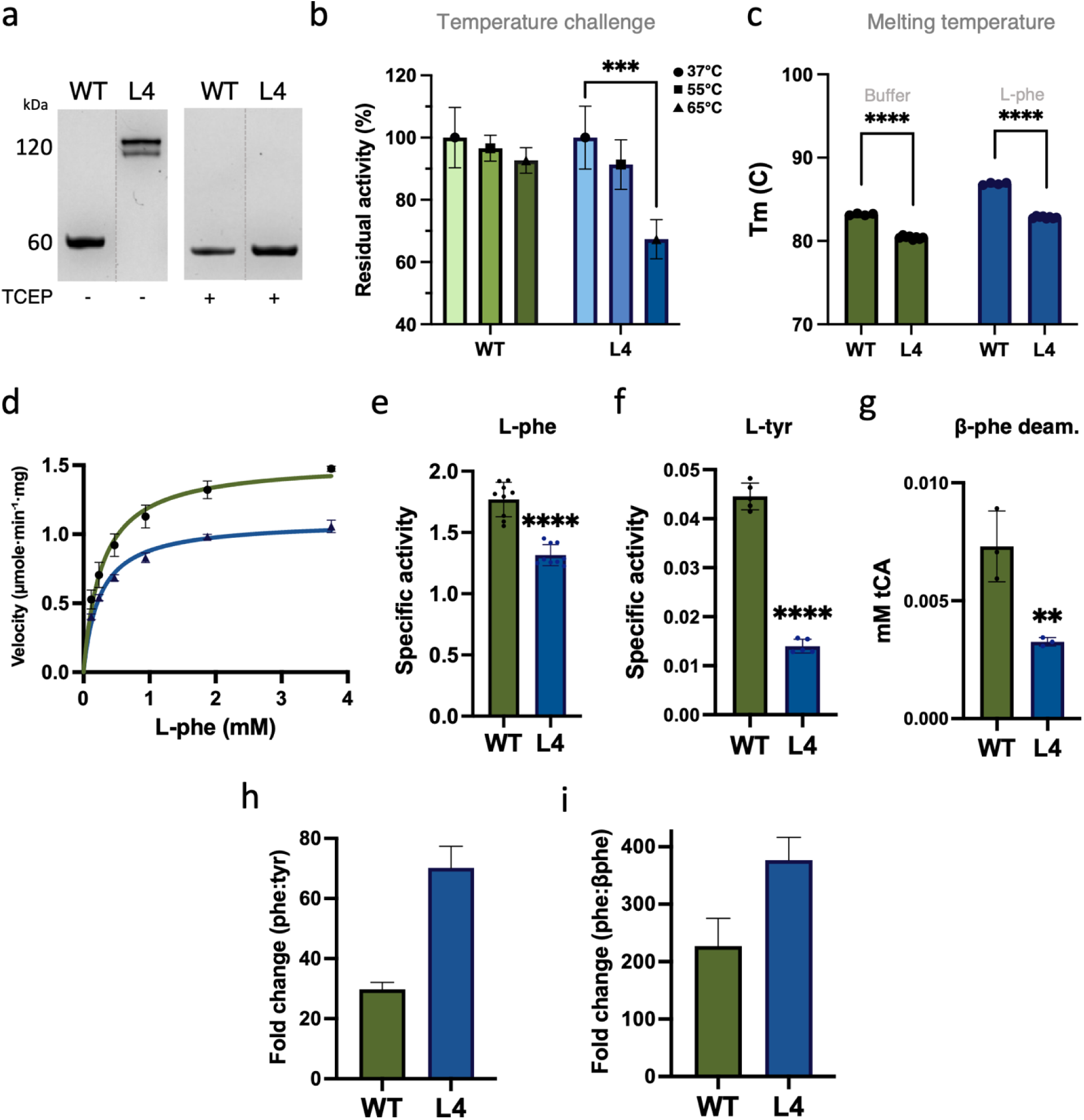
**Characterization of purified variant L4**. (a) SDS-PAGE of purified enzyme with and without TCEP (tris(2-carboxyethyl)phosphine hydrochloride). (b) Stability after one hour at 37, 55, or 65 °C. (c) Melting temperature in Tris-HCl buffer without and with saturating amounts of substrate. Data indicate loss of thermal stability of L4. (d) Michaelis-Menten kinetics for WT (green) and L4 (blue). Specific activity on (e) phenylalanine (30 mM), (f) tyrosine (2 mM), and (g) β-phenylalanine (30 mM) after 28 h of reaction time. Two-tailed unpaired t-test, * p < 0.05, ** p < 0.005, *** < 0.001, **** < 0.0001. Fold change calculated to show greater substrate specificity of L4 for phe over (h) tyr and (i) β-phe.

#### Disulfide bond at lid-like loop decreases the thermostability of L4

Disulfide bond engineering is primarily used for increasing protein stability. While our goal was not to improve stability, we assessed how the disulfide staple affected this phenotype. Interestingly, we found that L4 had reduced thermostability relative to WT, as determined by residual activity after a 1 h temperature challenge (**Figure 4b**). This destabilization was further supported by a Thermofluor assay, which revealed a decrease in melting temperature in the presence or absence of substrate (**Figure 4c**).

#### The L4 variant has enhanced substrate specificity

Michaelis-Menten kinetic analysis of WT and L4 reveals distinct differences in behavior after the addition of this disulfide bond. The K_M_ of L4 is reduced, reflecting enhanced binding “affinity” for phenylalanine (**Figure 4d**). While specificity increased, the v_max_ decreased ∼25 % (**Table 2**), suggesting that the disulfide bond enhanced substrate binding, yet imposes structural constraints that limit the conformational dynamics required for efficient turnover. PALs are well documented to have measurable catalytic activity on closely related aromatic amino acids such as L-tyrosine (tyr), yet phenylalanine remains the preferred substrate. Introduction of the disulfide bond at the base of the lid-like loop, variant L4, resulted in a reduction in phenylalanine turnover, with specific activity decreasing approximately ∼25 % (**Figure 4e**). Interestingly, the disulfide bond at the base of lid-like loop greatly reduced its activity on tyrosine (∼3×), indicating increased preference for phenylalanine (**Figure 4f,h**). Similarly, we found that β-phenylalanine deamination was also significantly decreased in L4 relative to WT (**Figure 4g**). Comparatively, L4 increases the specificity toward phe >2-fold relative to tyr (∼70× vs. ∼30×) and ∼1.7-fold relative to β-phe (∼375× vs. ∼225×) (**Figure 4h,i**).

**Table 2.**
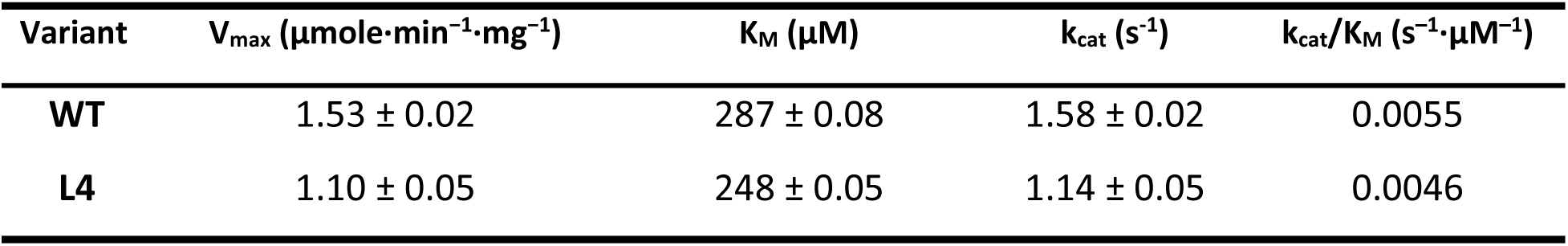
Kinetic constants of *Av*PAL variants.

#### Simulations indicate that L4 enables stabilization of phenylalanine in the active site

To evaluate and compare the binding affinity of phe and tyr, we used MM/GBSA (molecular mechanics/generalized Born surface area) calculations. The binding free energies (ΔG_bind) reflect configurational variability and indicate a clear substrate preference. Among the two substrates, phe exhibits the stronger binding energy at −31.9 ± 3.7 kcal/mol for L4, while tyr exhibits −21.2 ± 2.9 kcal/mol. The L4 variant also exhibits a substantially more favorable binding free energy than WT to phe (−31.9 ± 3.7 kcal/mol vs. −24.6 ± 2.6 kcal/mol), corresponding to an approximate ΔΔG of ∼7.3 kcal/mol in favor of the disulfide-bonded variant (**Figure 5a**). Tyr, in contrast, has a considerably smaller difference between WT (−19.8 ± 1.9 kcal/mol) and L4 (−21.2 ± 2.9 kcal/mol) at ∼1.4 kcal/mol, indicating only a non-significant change. These results demonstrate that the mutation selectively enhances binding affinity toward phe while having minimal energetic impact on tyr binding.

**Figure 5.**
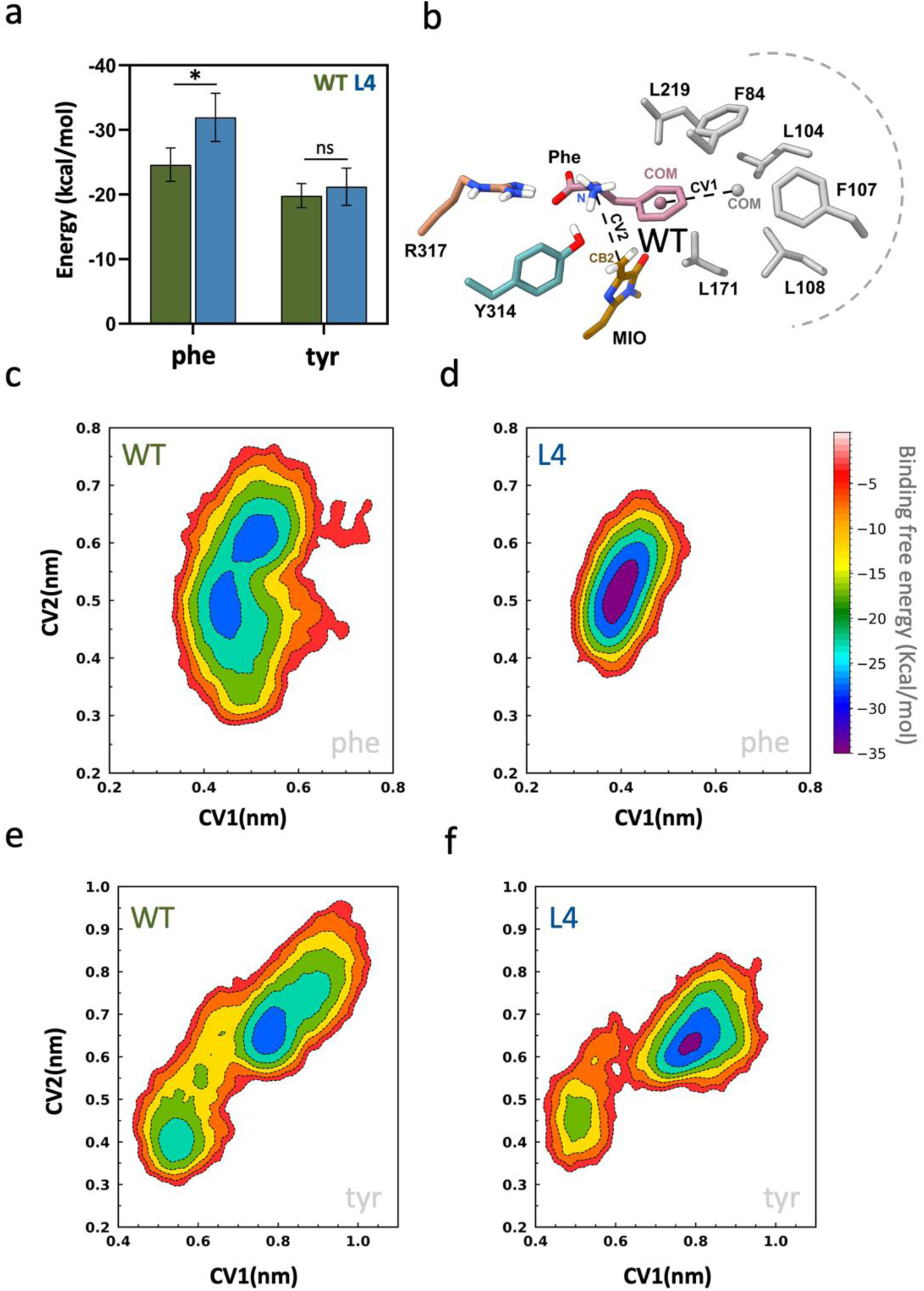
(a) Comparison of MM-PBSA binding free energies for phe and tyr between WT and L4. Bars represent the mean binding free energy (ΔG_bind, kcal/mol) calculated from MD trajectories, and error bars indicate standard deviation. Values are presented as mean ± SD (n = 3 per group). Statistical significance was assessed using an unpaired two-tailed t-test. \**p* < 0.05 was considered significant; ns, non-significant. (b) Structural representation of the active site, highlighting the defined collective variables (CVs): CV1 corresponds to the distance between the substrate aromatic ring COM and surrounding residues (F84, L104, F107, L108, L171, L219), CV2 corresponds to the distance between the MIO-Cβ2 atom and the substrate amino nitrogen. (c,d) Two-dimensional free-energy surfaces (FES) for phe in WT and L4; (e,f) FES for tyr in WT and L4.

Metadynamics simulations further confirmed the MM/GBSA binding energy results by exploring substrate conformations under applied bias potentials (**Figure 5**). As the only structural difference between the two substrates is the hydroxyl group on the aromatic ring, we specifically analyzed the interactions between the substrate’s aromatic ring and its surrounding pocket residues. To characterize these interaction dynamics, we defined two distance-based collective variables (CVs). CV1 was defined as the distance between the center of mass (COM) of the substrate aromatic ring and the COM of the surrounding pocket residues (F84, L104, F107, L108, L171, L219, and N223), which monitors substrate positioning and aromatic packing within the active site. CV2 was defined as the distance between the Cβ2 atom of the MIO moiety and the substrate nitrogen atom, representing a key reactive distance relevant to catalysis (**Figure 5b**).

The free energy surface analysis reveals a clear difference in how phe and tyr are accommodated within the active site. The WT displays a relatively broad free energy space, indicating conformational sampling in substrate positioning (**Figure 5c**). In L4, phe is confined to a single, narrow, and well-defined energy minimum, indicating stabilization in a dominant binding mode with limited conformational dispersion (**Figure 5d**). This minimum corresponds to favorable aromatic packing (CV1) within the hydrophobic pocket and optimal distance of the substrate amino group to the MIO reactive center. Although the global minima are centered within catalytically relevant distances (CV2), the broader energy surface for WT suggests dynamic sampling of multiple binding orientations.

Tyr exhibits broader conformational sampling along both CV1 and CV2 (**Figure 5e,f**). The presence of the hydroxyl substituent disrupts optimal aromatic packing for both WT and L4. This disruption leads tyr to explore a wider range of binding conformations within the active site. In CV2, tyr samples extended distances up to ∼9 Å, which shows that a significant portion of its conformations are non-catalytic. The hydroxyl group promotes adoption of non-catalytic geometries, reducing the likelihood of achieving a near-attack conformation. In contrast, phe exhibits a more confined free energy minimum in L4, maintaining favorable aromatic packing and optimal catalytic distances. Together with the MM/GBSA binding analysis, these observations indicate that L4 preferentially stabilizes phe in catalytically competent orientations, demonstrating higher substrate specificity toward phe compared to tyr.

#### QM/MM and QM/MM metadynamics reveal elevated energy barriers for L4

We performed QM/MM (quantum mechanics/molecular mechanics) and QM/MM metadynamics simulations to gain deeper insight into the catalytic mechanisms and electronic effects governing enzymatic function. These simulations integrate quantum mechanical (QM) treatment of the active site with molecular mechanics (MM) for the surrounding protein environment, enabling a more accurate representation of reaction pathways. The optimized structures of the key intermediates and transition states are shown in **Figure 6a**. The corresponding relative free energy profile (**Figure 6b**) shows the relative free energy differences along the reaction coordinate. The first transition state (TS1) in L4 has a higher energy barrier of 18.8 kcal/mol compared to WT’s barrier of 12.9 kcal/mol. The second transition state (TS2) shows a significant energy increase of 27.4 kcal/mol in L4 vs. 21.4 kcal/mol in WT, making the reaction pathway less thermodynamically favorable with L4. The contour plots (**Figure 6c,d**) show the free energy landscape along two CVs, where CV1 is the distance between the nitrogen atom on the substrate and the Cβ2 atom in the MIO moiety, and CV2 is the distance between the hydroxyl on tyr and the oxygen atom on the substrate. In WT *Av*PAL, the reaction pathway appears energetically smoother with distinct minima for IS1 and IS2, and shows a maximum energy barrier of ∼1.1 kcal/mol to form the intermediate states. The landscape for L4 is narrower, with increased energy barriers of ∼3.1 kcal/mol, suggesting increased conformational constraints and less efficient catalysis.

**Figure 6.**
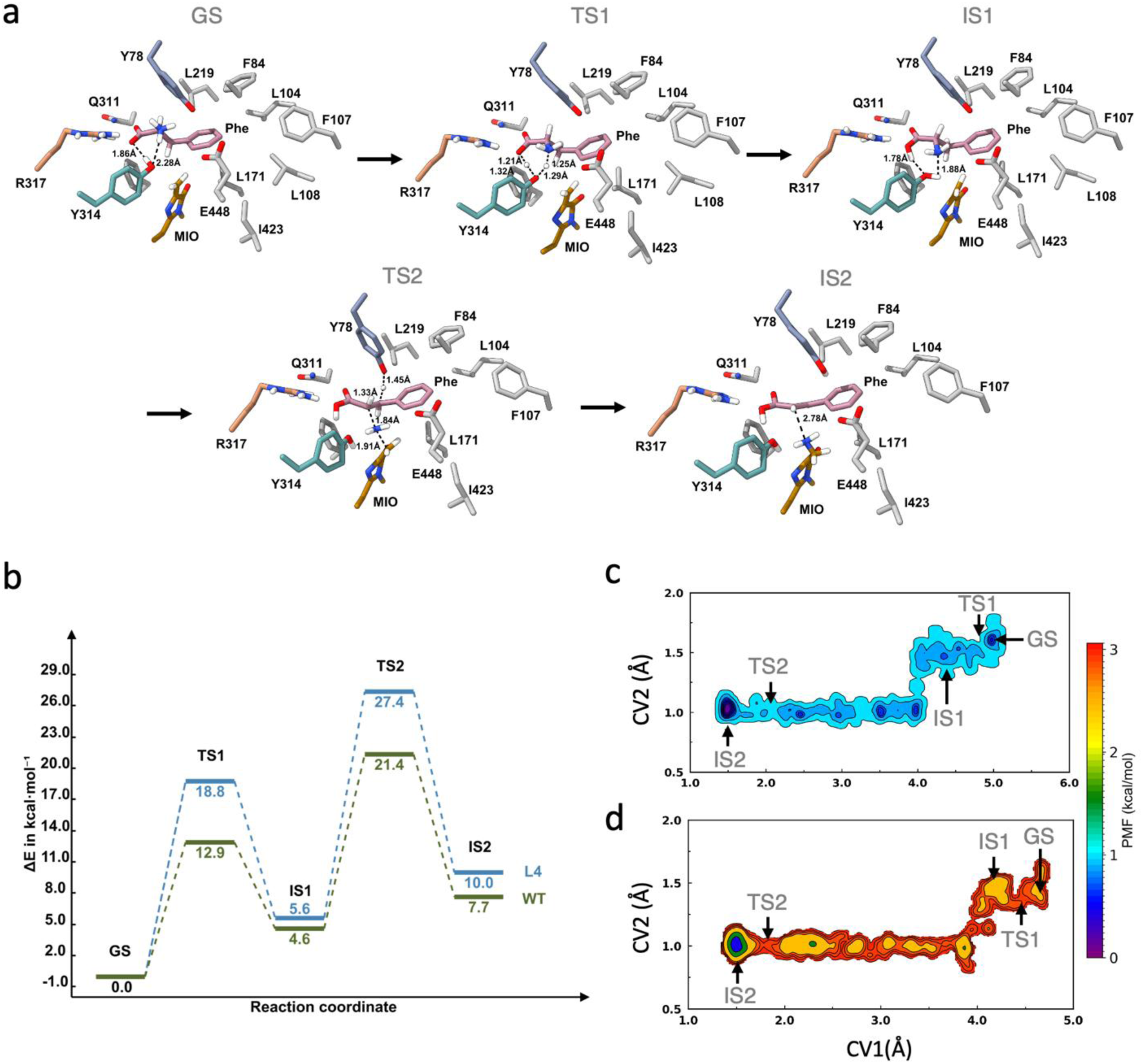
(a) Representative active-site configurations for the ground state (GS), transition states (TS1 and TS2), and intermediates (IS1 and IS2). Key active-site residues (Y78, Q311, R317, Y314, E448) and the MIO group are shown, highlighting their interactions with the substrate (phe). Only polar hydrogens are shown for clarity. All distances are in Å. (b) QM/MM free energy profiles along the reaction coordinate for WT and L4. (c) QM/MM metadynamics free energy landscape for WT projected along CV1 and CV2, showing a smooth reaction pathway with distinct ground (GS), intermediate (IS), and transition states; (d) Corresponding QM/MM metadynamics free energy landscape for L4, characterized by increased energy barriers. CV1 is the distance between the COM of the substrate and the COM of the backbone atoms of active site residues within 5 Å of the substrate; CV2 is the distance between the Cβ2 atom of the MIO moiety and the hydroxyl atom of Y78.

#### REMD reveals conformational rigidity of the lid-like loop in L4

To investigate the dynamic basis of the reduced activity in L4, we carried out REMD (replica exchange molecular dynamics) simulations. RMSF analysis revealed a pronounced reduction in flexibility within the lid-like residues. Whereas WT maintains substantial loop mobility that facilitates catalysis, L4 adopts a markedly rigid loop conformation. **Figure 7a** presents the average RMSF plot of the inner-lid loop, comparing flexibility across different REMD simulations. This rigidity likely restricts the conformational adaptability required for efficient substrate binding and turnover. To quantify the functional consequences of this rigidification, we monitored the distance between Y78, a key catalytic residue within the inner-lid loop, and the MIO moiety over 50 ns of REMD simulation. The WT exhibits pronounced fluctuations in the Y78-MIO distance, reflecting a dynamically adaptable lid loop that frequently samples open conformations required for efficient catalysis (**Figure 7b**). In contrast, L4 maintains a consistently shorter Y78-MIO distance with reduced fluctuation amplitude throughout the trajectory, indicating stabilization of closed or partially closed loop conformations (**Figure 7c**). Together, the RMSF and distance analyses demonstrate that the engineered disulfide bond in L4 imposes constraints on inner-lid loop motion. While this restriction enhances structural stability, it simultaneously limits the conformational flexibility required for productive catalytic turnover.

**Figure 7.**
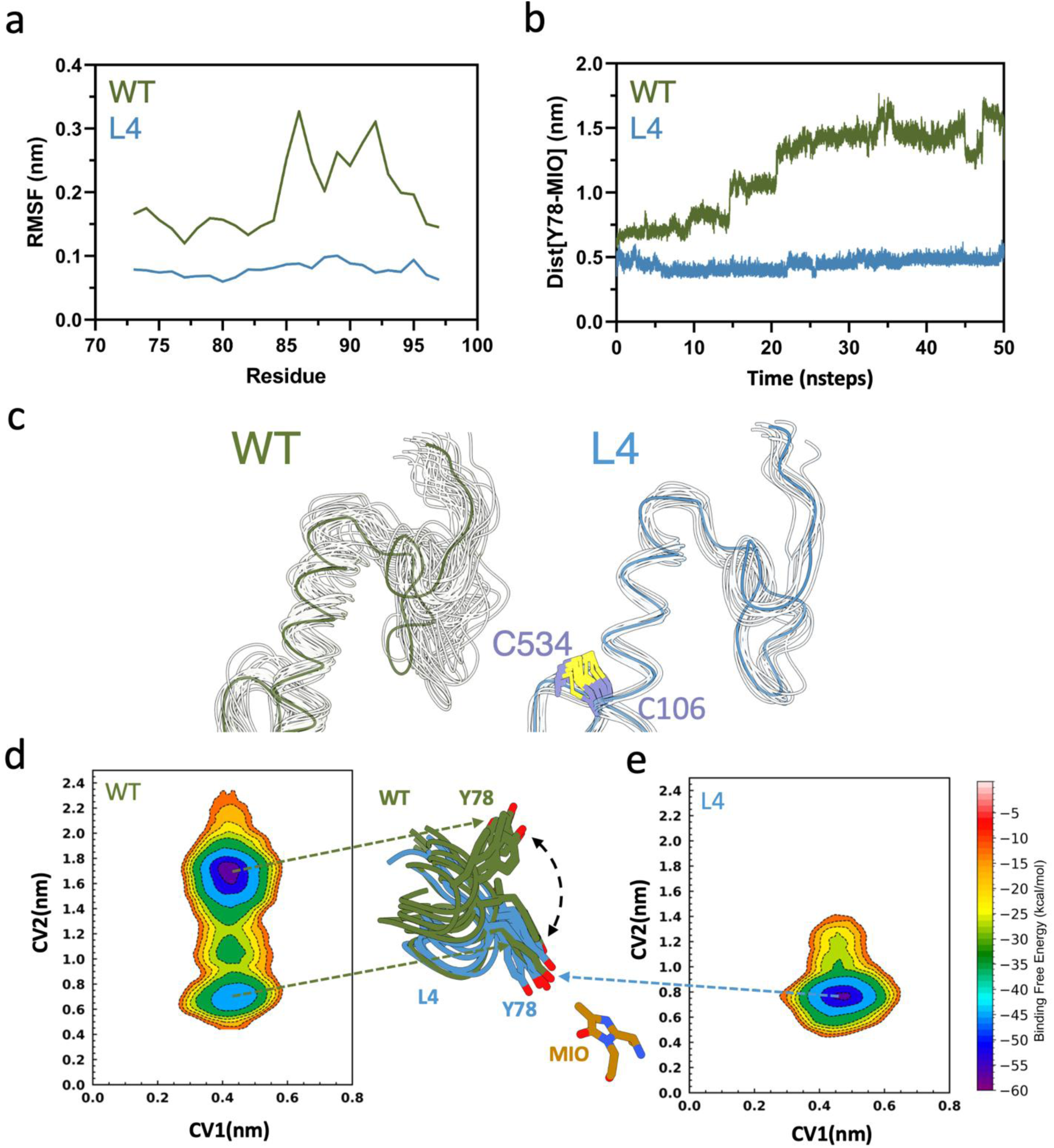
(a) Root mean square fluctuation (RMSF) profiles of the inner-lid loop region (residues 74–98) obtained from REMD simulations, comparing the WT enzyme (green) and L4 (blue). (b) The distance between Y78 and the MIO cofactor over 50 ns of REMD simulations averaged across multiple replicas, showing reduced fluctuations and a consistently shorter distance in L4 relative to the WT, indicative of restricted lid-loop motion. (c) Structural overlay of representative conformations highlighting the inner-lid loop region. (d,e) Two-dimensional free energy surface (FES) obtained from metadynamics simulations projected along collective variables CV1 and CV2, for WT and L4. The central structural overlay highlights differences around residue Y78 and the MIO group.

#### Metadynamics indicates restricted free energy landscapes in L4

To further characterize the free energy landscape associated with the substrate conformational space and active site, we employed metadynamics simulations. This enhanced sampling technique introduces a history-dependent biasing potential, allowing the system to overcome local free-energy minima and explore rare, yet functionally relevant, conformational states. Distance-based collective variables (CVs) are defined as follows: CV1 is defined as the distance between the COM of the substrate and the COM of the backbone atoms of active site residues within 5 Å of the substrate, and CV2 is defined as the distance between the Cβ2 atom of the MIO moiety and the hydroxyl atom of Y78. CV1 reflects changes in active-site dynamics relative to substrate positioning, whereas CV2 captures the flexibility of the in-lid loop. The residue Y78 functions as a key gating residue, undergoing opening and closing motions that regulate substrate access, as observed in the WT enzyme. Based on this structural insight, we explicitly incorporated Y78 into the CV definition to monitor loop dynamics during the simulations. The free energy surface (FES) plots obtained from the metadynamics simulations reveal distinct differences between WT and L4. CV2 exhibits two distinct energy minima (blue region) in WT, observed at approximately 0.6 nm and 1.8 nm (**Figure 7d**). This shows enhanced in-lid loop flexibility and broader conformational sampling. In contrast, L4 displays a more restricted conformational landscape, with a single minimum (blue region) centered around ∼0.8 nm (**Figure 7e**), reflecting more constrained and energetically limited in-lid loop dynamics.

#### Collective motion analysis confirms reduced functional dynamics in L4

To further examine the structural dynamics observed in the metadynamics and REMD simulations, principal component analysis (PCA) was applied to the simulation trajectories of both WT and L4. By decomposing the covariance matrix of atomic fluctuations, PCA identifies dominant modes of motion, providing insights into functionally relevant conformational changes. The porcupine plots (**Figure S1**) visualize significant motions along the first principal component (PC1), highlighting enzyme flexibility. In the wild-type (**Figure S1a**), pronounced flexibility is evident in the inner-lid loop region, with red spikes indicating the highest amplitude of motion, while green and blue regions signify moderate and minimal movement, respectively. Arrows depict correlated motion, emphasizing WT’s dynamic adaptability. In contrast, L4 (**Figure S1b**) exhibits a more constrained motion profile, with fewer high-amplitude fluctuations and dominant blue regions, indicating increased rigidity, particularly in the inner-lid loop.

## DISCUSSION

We report a strategy to selectively reduce the flexibility of a conserved lid-like loop in *Anabaena variabilis* (*Trichormus variabilis)* PAL through targeted disulfide bond engineering. We built and leveraged three complementary frameworks to predict cysteine residue pairs that introduce disulfide bonds. We found that the number of positions in this region was extremely limited, indicating that the enzyme provides a relatively narrow landscape for stabilizing cysteine pairs. Existing strategies to predict disulfide bonds are scarce, leading us to design our own multi-strategy cysteine prediction methodology. Our approach combined paired interaction energy (PIE) analysis, contact mapping, and machine-learning–based ranking to identify candidate disulfide-forming cysteine pairs in *Av*PAL. Despite this multi-pronged strategy, only a single engineered interchain variant, L4, formed a robust disulfide bond *in vivo* and *in vitro*. This outcome highlights a key limitation shared across disulfide prediction methods, that conformational dynamics are poorly captured by static models. While the geometry of a protein can be heavily inferred from its backbone, the functional behavior of a protein arises from the harmonic choreography of side-chain interactions that unfold across an ensemble of conformations rather than a single snapshot. Our results, therefore, support emerging views that successful disulfide engineering, particularly in flexible loops, requires explicit consideration of conformational ensembles rather than reliance on single structural assumptions.

Across variants, introduction of cysteine pairs reduced specific activity relative to WT, indicating that predicted geometrically permissible sites can introduce unfavorable steric or dynamic constraints on loop dynamics, active-site accessibility, or overall conformational flexibility. Notably, variants L1, L2, and L3 exhibited ∼50 % loss in activity, consistent with disruptions to loop mobility or altered access to the active site. Variant L4 displayed a comparatively moderate (∼25 %) reduction in activity, suggesting that certain positions may better tolerate cysteine substitution or stabilize conformations that remain catalytically competent. Disulfide bonds are commonly introduced to enhance protein thermostability, making the reduced stability of L4 unanticipated (10, 11). However, this loop has been repeatedly demonstrated to coordinate a crucial pivot point in the enzyme. This decrease in stability highlights the importance of accounting for full protein dynamics when attempting to engineer stability. Collectively, these findings highlight that flexibility can be stabilizing when it is functionally integrated into the protein’s energy landscape and that rigidity should not be assumed to correlate positively with thermostability. Our results indicate that geometric feasibility alone is insufficient for engineering novel disulfide bonds for the sake of improving global thermodynamic stability in highly flexible regions.

We demonstrate that significant rigidification of the lid-like loop resulted in a structural perturbation that modulates catalytic behavior by increasing the specificity towards L-phenylalanine, accompanied by reduced conformational mobility of both the catalytic residue, Y78, and the bound substrate within the active-site pocket. Intentional engineering of increased substrate specificity is mostly used in antibody engineering to increase specificity to a single target, while efforts in protein engineering are primarily focused on increasing catalytic efficiency (12). Alternatively, enzymes with minimal promiscuity are engineered to expand their capabilities or introduce completely novel specificities, i.e., enzymes with limited promiscuity are typically engineered to become more permissive, not less (13, 14). In this case, we believe that the reduction in conformations of positions in L4 enhances the specificity of phenylalanine due to better positioning for catalysis. The FES calculations demonstrate that L4 has an increased affinity for proton abstraction, supporting that better substrate positioning and reduction of energetic conformations have enhanced canonical specificity. One of the mutations in L4 is adjacent to residue F107, which has been reported to be involved in switching substrate preference from phe to tyr in aminomutases (4, 15). W106C in L4 also plays a role in substrate preference, albeit in the opposite direction – it further enhances phe preference. While unforeseen, our results highlight how dynamics of unstructured regions, like the lid-like loop of *Av*PAL, concurrently influence reaction rate (k_cat_), substrate specificity (phe vs. tyr vs. β-phe), and thermostability. Our work thus underscores the importance of studying unstructured loops in enzymes.

### EXPERIMENTAL PROCEDURES

#### Growth conditions, plasmids, and microbial strains

*Escherichia coli* strains used were DH5α (ThermoFisher, Waltham, MA), MG1655 *rph*^+^, BL21 (DE3) (ThermoFisher, Waltham, MA), and SHuffle Express (NEB, C3028J). Plasmid backbones used were pBAV1k (Addgene Plasmid #26702) and p15A (MilliporeSigma catalog #71147-3) (**Figure S2a**). Strains were plated using 1.5 % (w/v) agar and cultured in LB at 37 °C with rotary shaking at 250 rpm. Chloramphenicol was used at (25 µg/mL) whenever necessary. All culture media and chemical reagents were purchased from ThermoFisher Scientific (Waltham, MA), unless indicated.

#### Cloning and molecular biology

Site-directed mutagenesis PCR was performed to make cysteine substitutions. Primers were designed on Benchling and ordered from Genewiz. PCR amplification was done using Platinum SuperFi II (Invitrogen, Waltham, MA). Following PCR, a DpnI digestion was performed by adding 1 µL to each PCR reaction (up to 50 µL of PCR reaction volume) and incubating at 37 °C in a thermocycler for 1 h or overnight. PCRs were run on a 1 % agarose gel in TAE (tris-acetate-EDTA) buffer, followed by E.Z.N.A. gel extraction (Omega Bio-tek catalog #D2500-02) to collect fragments. Ligation was done with NEBuilder HiFi (NEB, E2621S). Ligated PCR products were chemically transformed into *E. coli* DH5α and verified by Sanger sequencing before cloning the final plasmids into *E. coli* MG1655 *rph*^+^. All enzymes, unless specifically noted, were obtained from New England Biolabs (NEB, Ipswich, MA).

#### PAL expression and protein purification

Double mutant *Av*PAL, C503S-C565S, was expressed in BL21 (DE3) using pBAV1k(1) (**Figure S2b,c**). *Av*PAL has an N-terminal 6x His-tag for IMAC purification. A 3 mL culture was set from one plated CFU and grown overnight at 37 °C before subculturing at 100× into 200 mL LB (Lysogeny Broth) supplemented with chloramphenicol. Cultures were grown for 16 h, then pelleted at 4500 ×g for 10 min at 4 °C. Pellets were saved in –80 °C for at least 24 h. *E. coli* SHuffle was used to purify all disulfide-bonded variants. A 3 mL culture was set from one plated colony and grown overnight at 37 °C before subculturing at 100× into 200 mL LB supplemented with chloramphenicol and grown overnight at 25 °C. For purification, 1.5 g of cell pellet was lysed using 10 mL of B-PER Reagent (ThermoFisher Scientific catalog #89821) and left to rock at 25 °C for 20 min. Lysed cells were aliquoted to 1.7 mL centrifuge tubes and spun at 20k ×g for 10 min at 4 °C to remove insoluble cell debris. Soluble supernatant was transferred to an IMAC resin gravity column. Approximately 500 µL of Ni-INDIGO resin (Cube Biotech catalog #75103) was used for each purification. Resin was equilibrated with two 5 CVs (column volumes) washes (equilibration buffer: 50 mM Na_2_HPO_4_, 300 mM NaCl, 10 mM imidazole), washed with five 5 CVs of wash 1 (50 mM Na_2_HPO_4_, 300 mM NaCl, 10 mM imidazole) two CVs of wash 2 (50 mM Na_2_HPO_4_, 300 mM NaCl, 20 mM imidazole), and eluted four times with 500 µL elution buffer (50 mM Na_2_HPO_4_, 300 mM NaCl, 200 mM imidazole). Elutions one and two contained the highest concentration of protein and were separately buffer exchanged into 25 mM Tris-HCl, pH 8.5, using 15K MWCO (molecular weight cut-off) Tube-O-DIALYZER (G-Biosciences catalog #786-613). Protein was quantified by Bradford method using BSA (bovine serum albumin) as the standard (ThermoFisher Scientific catalog #5561020) stored at 4 °C. Purity was measured with SDS-PAGE on NuPAGE Bis-Tris Mini Protein Gels, 4–12% (ThermoFisher Scientific catalog # NP0322BOX), imaged on ChemiDoc Go (Bio-Rad Laboratories, Waltham, MA).

#### Whole cell PAL activity assay

The whole cell assay was used to quickly measure tCA (*trans*-cinnamic acid) production by *Av*PAL and variants. A 3 mL culture of *E. coli* carrying one individual *Av*PAL variant plasmid was set to grow for 16 hours overnight at 37 °C. The next morning, 40 uL of the overnight culture was added into 4 mL of LB and antibiotic in a 24-deep well plate and grown to an OD of 0.4—0.5. The plate was spun down at 3000 ×g for 10 min and washed with PBS (phosphate-buffered saline) twice. The pellet was resuspended to a final volume of 1 mL. Washed cells were transferred to a 96-deep well plate for a final OD of 0.4 in 1 mL PBS pH 8.5 and 30 mM L-phenylalanine and incubated at 37 °C for 1 h. A 200 µL aliquot was removed to measure OD and tCA concentration. Final tCA production was determined by 290 nm/600 nm and normalized to WT, as described previously (5, 6).

#### Enzyme activity assay

The activity of *Av*PAL was measured by determining the rate of tCA production. 1 µg of purified protein was added to 200 µL of 30 mM L-phenylalanine in 50 mM Tris-HCl buffer, pH 8.5, in a UVStar 96-well plate (Greiner Bio-One catalog# 655801). The rate of tCA production was measured by reading absorbance at 290 nm every 10 s for 5 min at 37 °C using a SpectraMax M3 (Molecular Devices, San Jose, CA) plate reader, then quantified using a tCA standard curve to measure specific activity, as described previously (5, 6). Error bars represent SD. Uncertainty in ratios was calculated by propagation of error.

#### Stability assays: temperature challenge, melting temperature (thermofluor assay)

The temperature challenge assay was carried out by incubating purified protein at indicated temperatures (37, 55, 65 °C) in a thermocycler for 1 h, followed by 10 min at 37 °C before being transferred to a plate to measure specific activity. The melting temperature was determined by aliquoting ∼3 µg of purified protein with dye components (Protein Thermal Shift™ Dye Kit, Applied Biosystems). Proteins were added to qPCR plate and measured according to the dye kit specification. Melting curves were determined by Protein Thermal Shift™ (ThermoFisher Scientific)

#### SDS-PAGE and Western blot

To determine disulfide formation and protein purity, 1 µg of purified protein was added to 6× dye with and without 200 mM TCEP (tris(2-carboxyethyl)phosphine hydrochloride) and adjusted to a total volume of 16 µL. Samples were boiled at 70 °C for 10 min, then added to 4—12 % Bis-Tris gels. For downstream SDS-PAGE analysis, gels were washed three times for 5 min, then covered with SimplyBlue™ SafeStain (ThermoFisher Scientific catalog #LC6060) and left to develop overnight. The background was fully de-stained with water before imaging. For Western blot, the gel was transferred to a 0.45 μm nitrocellulose membrane (ThermoFisher Scientific catalog #88018) via wet tank transfer. Proteins were transferred for 1 h at 30 V surrounded by ice. The membrane was blocked in 3 % non-fat milk in 0.1 % TBST (ThermoFisher Scientific catalog # J77500.K8) for 1 h, washed three times for 5 min in 0.1 % TBST. Membrane was reacted with HRP-conjugated Mouse anti His-Tag mAb (ABclonal Technology catalog #AE028) for 1 h. Finally, the expression level of the target protein was detected by the enhanced chemiluminescence method via SuperSignal™ West Femto Maximum Sensitivity Substrate (ThermoFisher Scientific catalog #46640).

#### Pair interaction energy (PIE) determination

The PIE (e) between two amino acid residues refers to the energy associated with the non-covalent interactions, primarily, electrostatic and van der Waals interactions, when the residues are in proximity. The PIE for AvPAL was evaluated for all the residues, and the matrix of PIE was generated as described previously (16). To capture the dynamic interaction of residue interaction, short molecular dynamics (MD) simulations were performed at 303.15 K (see Molecular Dynamics (MD) Simulation section). The PIE between residues A and B was calculated using the Amber Param99, considering the electrostatic and van der Waals interactions over multiple simulation frames (17). The pairs of amino acids with the least and suboptimal pair interaction energies were visualized and validated as prominent hotspots for disulfide engineering. The formation of disulfide bonds among cysteine pairs and their stability were studied. The results can be found in the Supporting Data File.

#### Contact mapping

Contact maps were generated to analyze residue-level spatial proximity within *Av*PAL (18). Residue-residue contacts were defined based on the distance between selected atoms, typically the Cα or Cβ atoms of each residue (19). A residue pair was considered to be in contact if the interatomic distance was less than 5 Å. Contact maps were constructed on a residue-by-residue basis and used to identify candidate residue pairs for disulfide bond design. Contact frequencies were calculated over short MD simulations, which were performed at 303.15 K to capture the persistence of residue interactions (see Molecular Dynamics (MD) Simulation section). The contact score 𝐷 between residues A and B was calculated as the sum of the inverse of the contact (*f*) between atoms *x* from residue *A* and *y* from residue *B*, over multiple simulation frames *i* (**Equation 1**). Residue pairs with the highest contact scores were considered structurally optimal candidates and subsequently mutated to cysteine pairs for disulfide bond evaluation. The results can be found in the Supporting Data File.

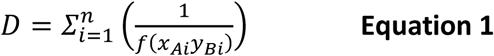

#### Machine learning

To further refine hotspot selection, a machine-learning framework was developed using curated datasets from PDBCYS (20), SPX (21), and an internally constructed dataset enriched with experimentally validated disulfide bonds. Short MD simulations were performed at 303.15 K for each protein to generate ensemble-based representations of cysteine pairs (see Molecular Dynamics (MD) Simulation section). For each protein ensemble, every cysteine-cysteine pair was characterized by 11 key descriptors over 100 simulation frames, including PIE (Δ*e*), contact score (Δ*D*), and geometrical parameters governing disulfide bond formation. Here, Δ represents the change between simulation frames and the initial/static structure. These parameters collectively capture the energetic and packing stability of the cysteine environment.

Geometrical descriptors included:

● C: the distance between the Cα atoms of two cysteine (cys) residues,
● d: the distance between the Sγ atoms of two cys residues,
● α_1_: the angle formed between Cα-Cβ-Sγ,
● α_1_: the angle formed between Sγ- Sγ’-Cβ’,

χ: the torsional angles formed between the planes of the two cys residues, including χ**_1_** (N-Cα-Cβ-Sγ), χ**_2_** (Cα-Cβ-Sγ-Sγ’), χ**_3_** (Cβ-Sγ-Sγ’-Cβ’), χ**_2’_** (Sγ-Sγ’-Cβ’- Cα’), and χ**_1’_** (Sγ’-Cβ’- Cα’-N’) The final descriptor matrix (11×100) was used as input for a convolutional neural network (CNN) designed to learn both local and global patterns. The first convolutional layer, with 32 filters of size 3×3, captured early correlations. A Rectified Linear Unit (ReLU) activation function introduced non-linearity, followed by a max-pooling layer to reduce dimensionality while preserving essential features. Deeper convolutional layers with 64 and 128 filters extracted higher-order relationships between dynamic geometries and energetics. Flattened outputs were passed through dense layers for feature integration, and a final sigmoid layer predicted the probability of disulfide bond formation. The model was trained using binary cross-entropy loss and validated on unseen structures.

#### Molecular dynamics (MD) simulation

All predicted disulfide variants from the three computational approaches were modeled using AlphaFold (22), and the cofactor MIO was positioned according to the reference crystal structure (PDB 2NYN). MD simulations for all variants were performed using the AMBER99SB force field (23) within the GROMACS 2023 package (24). The protein complex was placed in a triclinic box and solvated with SPC216 water molecules, extending 10 Å beyond the protein in all directions (25). The system was neutralized, and an ionic strength of 0.15 M was maintained by adding the appropriate number of Na⁺ and Cl⁻ counterions. Energy minimization was carried out using the steepest descent method (26). All MD simulations were performed with a 2fs integration time step. The minimized systems were equilibrated under the canonical (NVT) and isothermal-isobaric (NPT) ensembles for 1 ns each at 300 K. Production MD simulations were then carried out across a temperature range (303.15 K,313.15 K, 323.15 K, 333.15 K, 343.15 K, and 353.15 K), with temperature increments applied every 3 ns, for about 20 ns. Trajectory analyses were performed using the GROMACS package, including calculations of root mean square deviation (RMSD), root mean square fluctuation (RMSF), and sulfur-sulfur distances between cysteine residues in the variants using gmx mindist (24).

#### Replica exchange molecular dynamics (REMD)

REMD simulations (27) were performed using GROMACS 2023 compiled with gmx_mpi, using AMBER99SB force field. System preparation, including structure optimization, solvation, ionization, energy minimization, and equilibration, followed the same protocol as for conventional MD simulations, which is detailed in the previous section. For the production runs, 14 replicas of the system were simulated in parallel with temperatures exponentially distributed between 303.15 K and 373.15 K, totaling 700 ns (50 ns, each replica) of simulation time. Exchanges between neighboring replicas were attempted periodically at every 500 ps, with temperature gaps adjusted to maintain an approximately constant exchange probability. Temperature was controlled using Langevin dynamics with a collision frequency (γ) of 1 ps⁻¹. Bond vibrations involving hydrogen atoms were constrained using the SHAKE algorithm, allowing a larger integration time step (28).

#### Metadynamics simulations

Well-tempered metadynamics simulations (29) were performed using the PLUMED library (30), starting from structures obtained from NPT equilibration. These simulations were employed to evaluate the effects of mutations on substrate conformations and active-site dynamics. The collective variables (CVs) were carefully selected to capture mutation-induced conformational rearrangements of the substrate and the active site, as the appropriate choice of CVs is critical for accurately characterizing changes in active-site flexibility and their impact on catalytic efficiency. Both the wild-type and variant L4 were simulated for 50 ns, using seeds for initial velocity generation. The metadynamics parameters were set as follows: PACE = 500, HEIGHT = 1.0 kJ/mol, SIGMA = 0.01, bias factor = 2, and temperature = 310 K.

#### QM/MM simulations

Representative equilibrated structures of *Av*PAL wild and variant from MD simulations were subjected to hybrid quantum mechanics/molecular mechanics (QM/MM) simulations to probe the impact of disulfide engineering on the catalytic reaction environment. The enzyme-substrate complexes were solvated in a cubic water box with a minimum solute-solvent separation of 12 Å in all directions using the QwikMD interface implemented in VMD. System electroneutrality was maintained by adding NaCl to a final concentration of 0.15 M. Protein parameters were assigned using the CHARMM36 force field (31), while TIP3P water molecules were used to model the solvent environment. Non-bonded interactions were treated using a 12.0 Å cutoff for short-range interactions, and long-range electrostatics were handled using the particle-mesh Ewald (PME) method. Equations of motion were integrated using the r-RESPA multiple time-step algorithm, with a 2fs integration step. Initial system relaxation involved 1,000 steps of energy minimization using the conjugate gradient method, followed by thermal equilibration at 300 K under NPT conditions using a Langevin thermostat (collision coefficient of 1 ps⁻¹) and a pressure of 1 atm.

Following classical equilibration, QM/MM partitioning was performed using the QwikMD QM/MM interface. The QM region encompassed the MIO adduct, the substrate, and key catalytic residues Y314 and R317, along with nearby water molecules within 3.5 Å of the reactive center. The total charge of the QM region was constrained between +1 and −1 to ensure numerical stability during semi-empirical calculations (32). The QM/MM system was further minimized for 1,000 steps and equilibrated using 10,000 steps of simulated annealing. Production QM/MM simulations were carried out using the PM7 Hamiltonian for the QM region in conjunction with the CHARMM36 force field for the MM region.

#### QM/MM metadynamics

To characterize the free energy landscape of the catalytic step and assess the influence of disulfide-induced structural stabilization on reaction energetics, QM/MM metadynamics simulations were performed. The equilibrated QM/MM structure was used as the starting configuration. Simulations were conducted at 300 K and 1 bar under periodic boundary conditions using NAMD (33) with the colvars module (34). Metadynamics simulations employed a 0.5 fs integration time step and were run for 1,000 ps. Two chemically relevant collective variables (CVs) describing the progression of the reaction coordinate were selected, corresponding to distances involving key reactive atoms in the substrate, MIO adduct, and catalytic residues. Bias potentials were constructed by depositing Gaussian functions with a height of 0.2 kcal/mol at regular intervals along the CVs, enabling enhanced sampling and escape from local minima. The QM region was treated using the PM7-based QM/MM framework throughout the metadynamics simulations. The resulting free energy profiles were used to compare reaction feasibility and energetic barriers between the wild-type enzyme and the variant.

#### Principal component analysis (PCA)

PCA (35) was employed to characterize the dominant collective motions of *AvPAL* and to assess the impact of disulfide engineering on protein conformational dynamics. PCA was performed on the Cα atomic coordinates extracted from the equilibrated portions of the MD trajectories for both the wild-type enzyme and Variant L4. Prior to analysis, overall translational and rotational motions were removed by least-squares fitting each trajectory frame to a reference structure.

The covariance matrix of positional fluctuations was constructed using mass-weighted atomic displacements according to **Equation 2**:

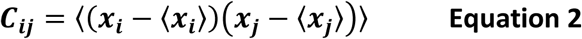

where 𝑥_𝑖_and 𝑥_𝑗_represent the Cartesian coordinates of atom 𝑖and 𝑗, respectively, and angular brackets denote time averaging over the trajectory. Diagonalization of the covariance matrix yielded eigenvalues and eigenvectors corresponding to the amplitude and direction of the principal motions. The first few principal components (PCs), accounting for the majority of the total variance, were selected for further analysis. The extent and direction of the most dominant motions of all complexes were visualized through porcupine plots using visual molecular dynamics (VMD) software.

## DATA AVAILABILITY

All data are contained within the manuscript.

## SUPPORTING INFORMATION

This article contains supporting information.

## ACKNOWLDGMENTS

We thank Dr. Todd C. Chapell (Enrich Bio, Waltham, MA), Dr. Vikas D. Trivedi (St. Jude Children’s Hospital, Memphis, TN), Prof. Ekaterina Heldwein (Tufts University, Boston, MA), Prof. James Baleja (Tufts University, Boston, MA), Dr. Cynthia Collins (Ginkgo Bioworks), and members of the Nair lab for their helpful comments and suggestions.

## AUTHOR CONTRIBUTIONS

Rebecca Condruti: Formal analysis, Investigation, Methodology, Visualization, Writing – original draft, Writing – review & editing Likith Muthuraj: Formal analysis, Investigation, Methodology, Visualization, Writing – original draft Jeevan K. Prakash: Formal analysis, Investigation, Methodology, Visualization, Writing – original draft Samuel D. Littman: Investigation Pravin Kumar R.: Conceptualization of computational methods, Supervision, Project administration, Funding acquisition, Writing – review & editing Nikhil U. Nair: Conceptualization, Supervision, Project administration, Funding acquisition, Writing – review & editing

## FUNDING AND ADDITIONAL INFORMATION

This work was supported by National Institutes of Health award R21HD105934 (to N.U.N. & P.K.R.), R01AG069917 (to N.U.N.), and National Science Foundation award CBET-2208390 (to N.U.N. & P.K.R.).

## SUPPORTING INFORMATION

This article contains supporting information.

