## Supplemental Information for "An engineered disulfide staple restricts lid loop dynamics and alters substrate specificity of phenylalanine ammonia-lyase"

### RUNNING TITLE

Disulfide Staple Alters Substrate Specificity of PAL

Figure List:

1. Supplemental Figure 1: Principal Component Analysis (PCA) of enzyme dynamics.
2. Supplemental Figure 2: Description of expression conditions for *Av*PAL.

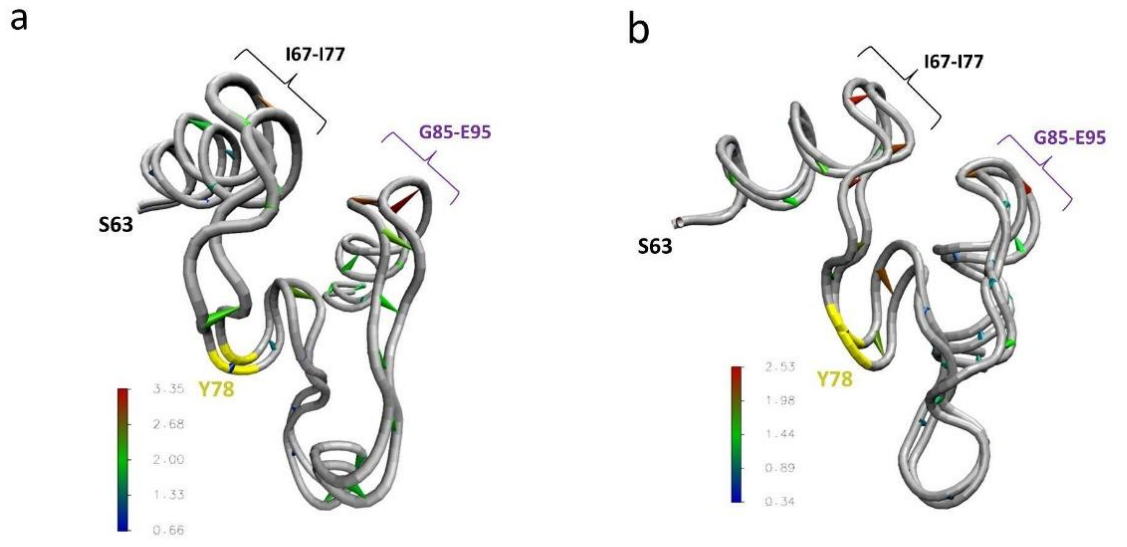


**Figure S1.** Principal Component Analysis (PCA) of enzyme dynamics.
(a) Porcupine plot for the WT enzyme from metadynamics, showing significant motion in the inner-lid loop. (b) Porcupine plot for L4 from metadynamics, illustrating a more constrained motion profile with reduced high-amplitude fluctuations, indicating increased rigidity in the inner-lid loop. Red indicates the highest movement, followed by green (moderate) and blue (least). Arrows represent the direction of correlated motion.

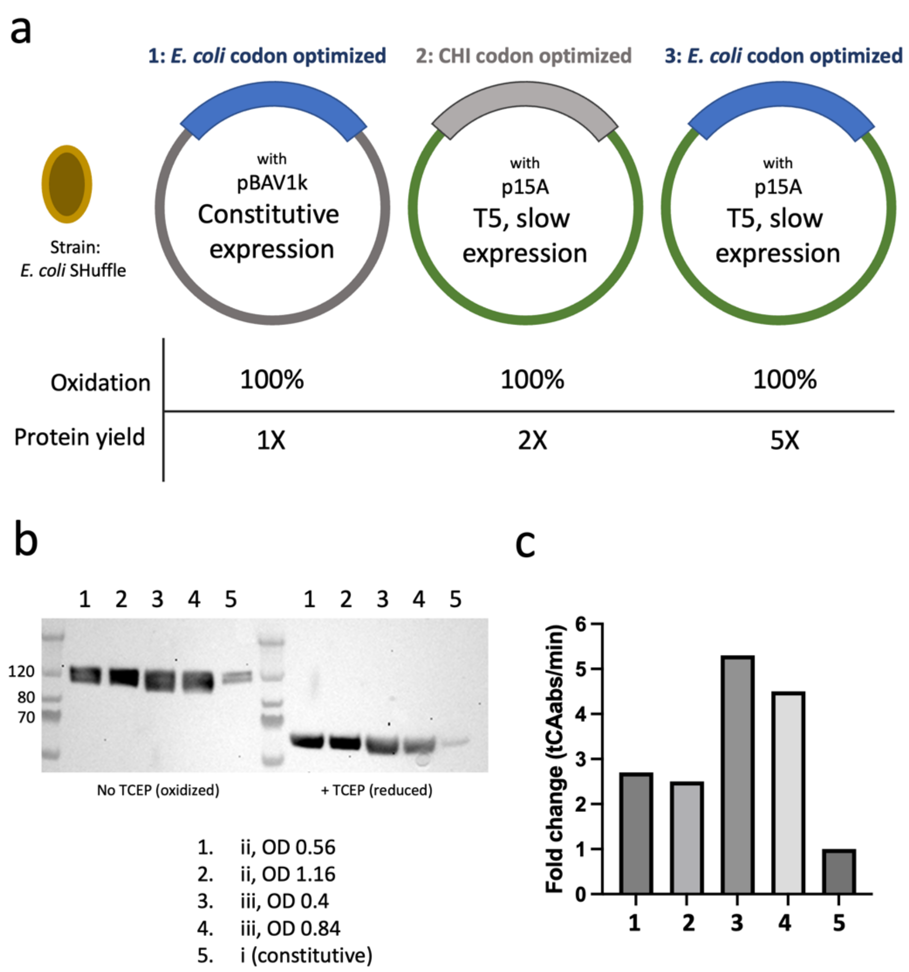


**Figure S2.** Description of expression conditions for *Av*PAL.
(a) Cartoon representation of paired codon optimization strategy with vector expression (1). Protein yield was best with basic codon optimization and T5 inducible expression as compared to previously published methods (2). (b) Western blot of protein yield from lysate, normalized to total lysate concentration. (c) Fold change of enzyme activity from lysate, normalized to total lysate.
